# The complex microscopic dynamics of cells in high-density epithelial tissues

**DOI:** 10.1101/2025.09.04.674282

**Authors:** Yuan Shen, Wang Xi, René-Marc Mège, Walter Kob, Benoit Ladoux

## Abstract

Epithelial tissues line the surfaces of vital organs and are often densely packed in a disordered, mechanically arrested state, referred to as ‘jammed’ or ‘glassy state. While collective migration at low-density regimes has been extensively studied, the microscopic dynamics within such jammed epithelia remains poorly understood. Here, we reveal that contrary to expectations from thermal systems with glassy dynamics, jammed epithelial monolayers do not exhibit cage effects that reflect the temporary spatial trapping of the particles. Instead, cells display sub-diffusive creep and Fickian yet non-Gaussian dynamics, accompanied by compressed exponential relaxation, features that reveal stress-driven fluidity. We show that cell divisions and extrusions transiently do enhance local motion, they are insufficient to fluidize the tissue globally. Fast-moving cells form collective, anisotropic clusters, and these dynamic heterogeneities correlate with local structural entropy and low-frequency vibrational modes. These findings challenge the conventional view that jammed tissues are static and inert structures, uncovering a hidden fluidity that can be expected to play a critical role in morphogenesis, wound healing, and early tumor progression.

## Introduction

Epithelial tissues are essential for maintaining organ integrity, guiding morphogenesis, and repairing wounds. These monolayers often experience dynamic restructuring (*1*), yet in many physiological settings, such as mature barriers or early-stage tumors, they are packed densely and disorderedly and appear mechanically arrested (*2, 3*). This so-called “jammed” state (*4, 5*), in which cells are unable to move freely, has been compared to glasses (*6, 7*). Understanding how individual cells move within such dense tissues is crucial for elucidating how epithelial layers retain plasticity during biological processes such as morphogenesis, regeneration, and cancer progression (*8*).

At low densities, epithelial cells exhibit collective migration, forming turbulent flows or flocking patterns that have been extensively characterized (*9-12*). In contrast, high-density monolayers exhibit strongly reduced mobility and enhanced mechanical rigidity — features commonly associated with a jamming transition (*4*). This rapid slowing down of cell dynamics is often described using analogies to glass-forming materials, in which particles become trapped in “cages” formed by their neighbors (*13*). Inspired by this analogy, several theoretical models have predicted that epithelial cells in the jammed state should exhibit local confinement and slow, glassy dynamics (*14-22*).

However, the extent to which these models capture the true behavior of living tissues is an open question. Experimentally, direct measurements of single-cell dynamics in high-density monolayers are rare (*23*), and previous observations of caging have often been inferred rather than quantified (*23-25*). Moreover, theoretical studies suggest that active processes—such as cell division and extrusion—can promote tissue fluidization even at high densities, thereby challenging the notion that densely packed tissues behave like glassy materials (*26, 27*).

In this work, we present a comprehensive experimental investigation of the microscopic dynamics of epithelial cells in the jammed state. By tracking thousands of cells over long-time scales, we demonstrate that high-density epithelia do not exhibit glass-like caging, but instead display creep-like motion, dynamic heterogeneity, and fluid-like relaxation. We further reveal that rearrangement-prone regions show significant correlations with structural and vibrational properties, suggesting that jammed tissues do retain mechanical susceptibility that enables localized relaxation. These findings not only challenge prevailing assumptions about epithelial glass, but also provide a new framework for understanding how tissues balance stability with dynamic responsiveness.

## Results

### Experimental Setup

MDCK cells stably expressing Histone1-GFP are cultured at high density (∼9,400 cells/mm^2^) on glass substrates coated with polymerized Matrigel. The samples are cultured for 2 weeks before imaging to ensure strong jamming. Cell nuclei are tracked using deep learning-based algorithms, achieving a tracking accuracy of approximately 100 nm (see Fig. S1 and *Methods* for details). Figure 1 shows a snapshot of the cell nuclei revealing that on large length scales the structure is disordered while for short distances the packing shows a liquid-like arrangement. The inset quantifies this local order by means of the pair correlation function of nuclei, g(**r**), calculated in a local coordinate system that is aligned with the long and short axis of the cells, defining, respectively, the *x*’ and *y*’ axes. The location of the first peak in g(**r**) is about 13 μm along the *x*’-axis and 10 μm along the *y*’-axis, respectively. The function in the non-aligned lab coordinate system can be found in Fig. S2, and displays the typical isotropic ring pattern found in disordered systems ^10^.

**Fig. 1.**
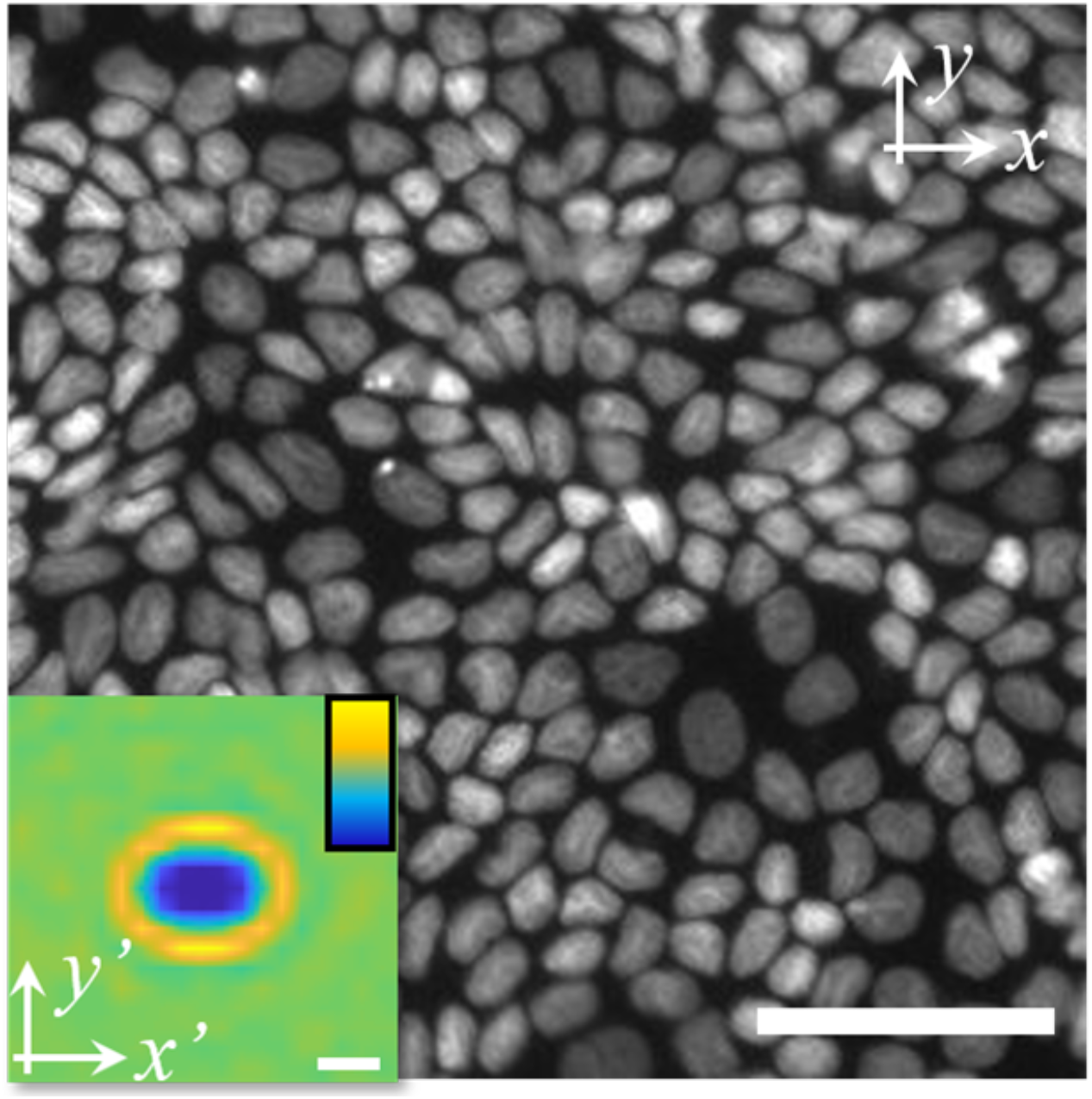
Spatial arrangement of the cells. Microscopy image of nuclei of MDCK cells at high density. Scale bar is 50 μm. Inset: Pair correlation function of nuclei g(**r**) in a local coordinate frame with the long-axis of the reference cell nucleus being aligned along the *x*’-axis. Scale bar is 10 μm. The color varies linearly from 0 (dark blue) to 1.5 (light yellow).

### Absence of cage effects revealed by the mean square displacements

In systems exhibiting cage effects, particles typically remain confined within small regions for extended periods, occasionally making large jumps—often exceeding the particle size—as they hop to a new cage ^10^. However, no such behavior is observed in the nuclei motion as can be concluded from Figure 2a which shows representative trajectories of cell nuclei. Instead, the trajectories appear to be continuous, lacking the large, abrupt hops characteristic of cage-breaking events.

**Fig. 2.**
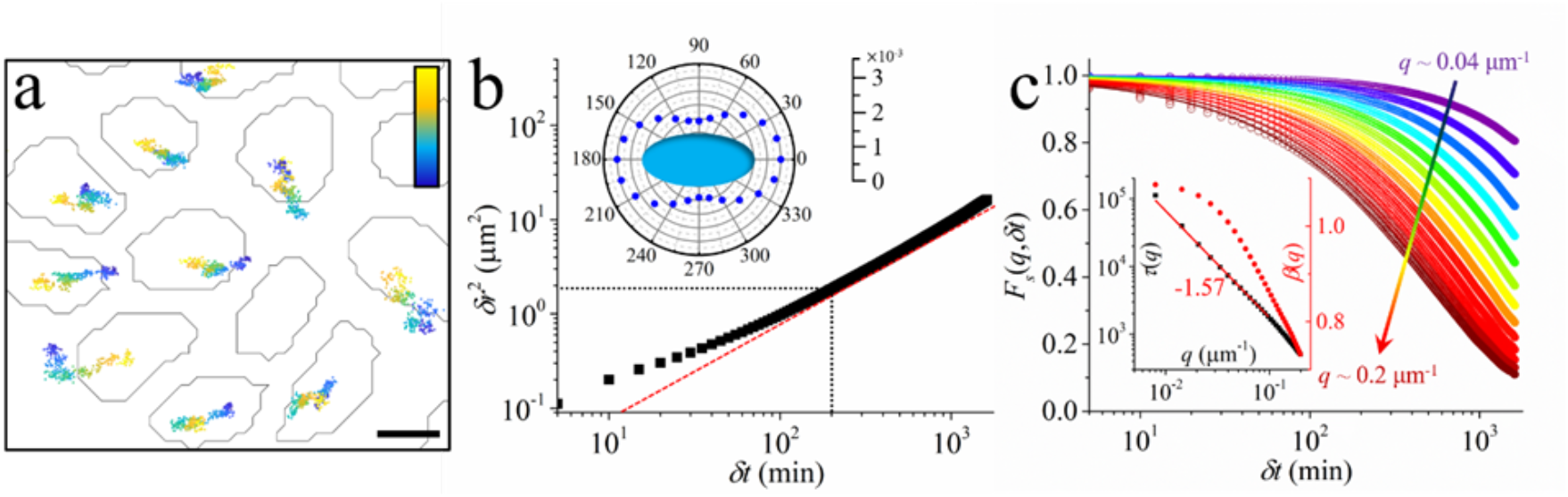
Microscopic dynamics of cells. (a) Trajectories of cells. Scale bar 5 μm. The color bar scales linearly from dark blue to light yellow, which represents the time lapse from 0 to 1625 min. (b) Translational mean squared displacement of cells as a function of time delay. The dashed black lines represent the cross-over from sub-diffusive regime to normal diffusive regime. The red dashed line represents a power-law with an exponent of 1. Inset: The diffusion coefficient of cells along different directions with respect to the long axes of the cell nuclei (unit: μm^2^ min^-1^). (c) Incoherent intermediate scattering function for different values of the wave vector **q** which increases from 0.04 to 0.2 μm^-1^ in a step of ∼0.006 μm^-1^ as indicated by the arrow. The solid lines are fits of the form 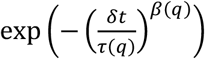. The inset shows *τ*(*q*) and *β*(*q*) as a function of *q*. The peak in the static structure factor is around *q* = 0.1 μm^-1^.

To determine quantitatively if there is a cage effect, we probe the translational mean squared displacement (TMSD) (Fig. 2b, *Methods*). For systems exhibiting cage effects, the TMSD shows a pronounced plateau at intermediate times, reflecting transient confinement that persists over a broad range of time scales (*13, 28, 29*). However, unlike the predictions of some vertex and Voronoi models (*16-22*), our data does not show such a plateau. To eliminate the possible influence of local collective cell migration related to Mermin-Wagner fluctuations, we have calculated the “cage related TMSD”, Fig. S3a (*Methods*), and also here no evidence of cage effects is found. Instead, the TMSD exhibits robust sub-diffusion at small *δt* and diffusive growth at large *δt*, confirming that here the cage effect observed in thermal glassy systems is absent.

It is noticed that the overall TMSD reaches ≈20 µm^2^ (Fig. 2b), i.e. a root mean square (RMS) displacement of ≈4.5 µm. Could this be explained by nuclei wobbling inside cells? At our confluence (≈ 9,400 cells mm^−2^), the mean lateral cell footprint is ≈ 106 µm^2^ (equivalent circular diameter ≈ 11.6 µm). With nuclear diameters of order 8−10 µm, simple geometry bounds any intra-cell nucleus-centroid shift to ≤ (*D*_cell_−*D*_nucleus_)/2 ≈ 1-2 µm before contacting the cortex. This 1−2 µm is a conservative geometric ceiling; cytoskeletal tethering and cytoplasmic viscoelasticity make typical intra-cell nuclear fluctuations substantially smaller, whereas we measure ∼4.5 µm RMS— well beyond intra-cell wobble and thus indicative of tissue-level motion. Consistently, (i) the TMSD shows no long-time plateau expected for a nucleus caged in a cell (Ornstein−Uhlenbeck) (Fig. 2b), (ii) the incoherent intermediate scattering function exhibits compressed relaxation at small *q* (Fig. 2c), (iii) divisions/extrusions induce transient flow over ∼20 µm (Fig. 3), (iv) short-time steps display directional anti-correlations (creep-like memory; Fig. 4a), (v) velocity–velocity correlations extend to ∼20 µm (Fig. 5g), and (vi) rearrangements correlate with structural entropy and soft vibrational modes (Fig. 6) — all signatures of mechanically coupled, tissue-level dynamics rather than independent nuclear jiggle.

**Fig. 3.**
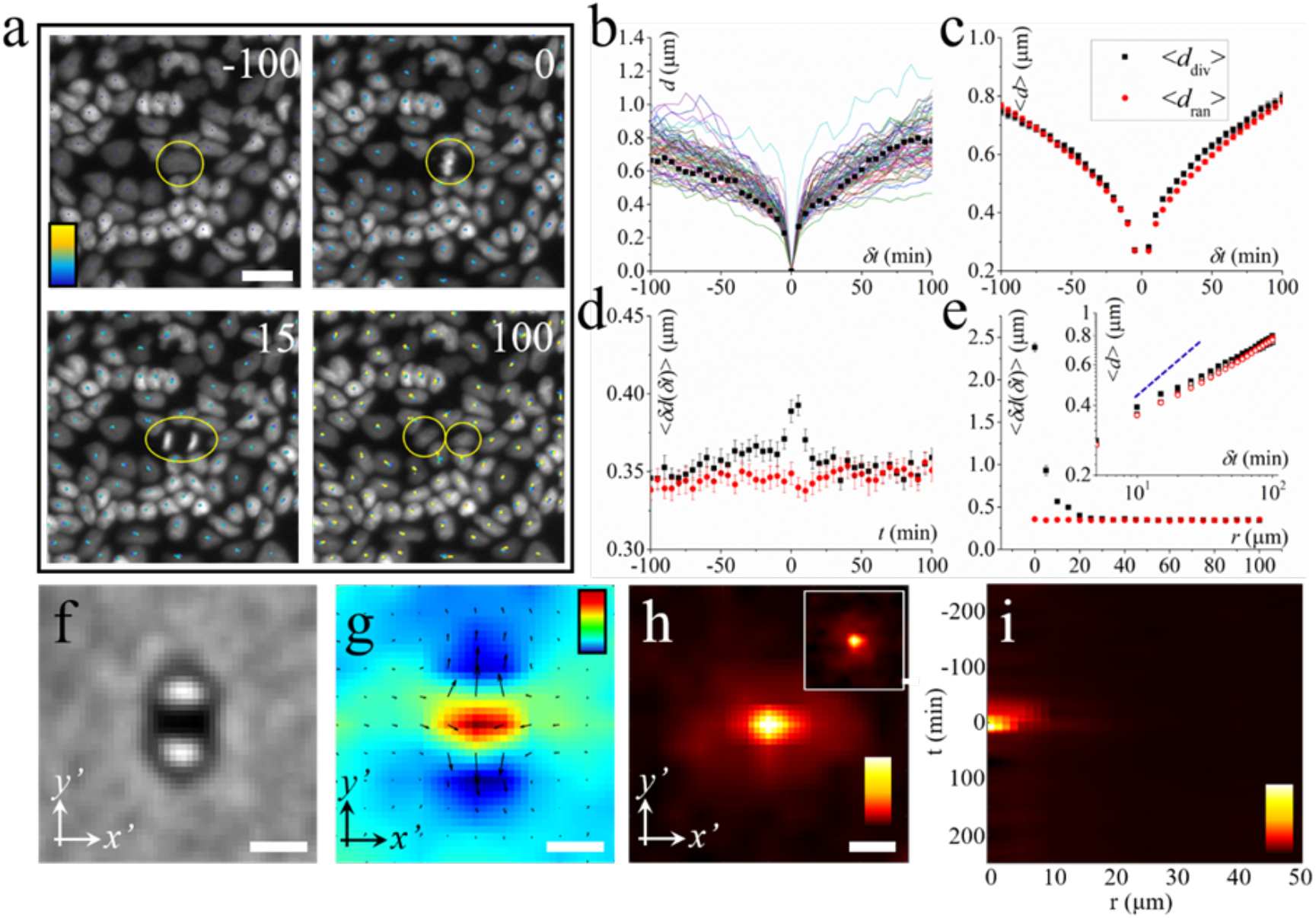
Dynamics induced by single cell divisions. (a) Snapshots showing the division of a cell at different moments. The dots represent cell trajectories. The color bar represents time lapse which scales linearly from 0 min (dark blue) to 200 min (light yellow). The dividing cell is marked with a yellow circle. Scale bar is 20 μm. (b) Cell displacement, *d*, averaged over local cells around a dividing cell as a function of time. *δt* represents time delay before and after division. Solid lines of different colors represent cell displacements of different division events. The black squares represent the cell displacement of the division event in (a). (c) Mean cell displacement, <*d*>, averaged over different division events (black squares, <*d*_div_>) as a function of time. The red dots represent the mean cell displacement averaged over different random control regions, <*d*_ran_>. (d) Mean displacement step, ⟨*δd*(*δt*)⟩, of local cells averaged over different division events (black squares) as a function of time before and after division moment. The red dots represent ⟨*δd*(*δt*)⟩ of local cells in random square regions. (e) ⟨*δd*(*δt*)⟩ at the division moment as a function of distance away from the division event. The black (red) dots represent ⟨*δd*(*δt*)⟩ of local cells in division (random) square regions. The inset shows the loglog plot of *d*_*div*_(*δt*) (black) and *d*_*ran*_(*δt*) (red) in (c) as a function of time. The solid (hollow) symbols represent the mean cell displacements after (before) division. The blue dashed line indicates a power law function with an exponent of 1/2. (f) Microscopic image obtained by averaging the brightfield images of different division events. Each brightfield image is rotated so that the long axis of the condensed chromosome is aligned along the *x*’-axis. Scale bar 10 μm (g) Mean velocity field obtained by averaging the PIV (particle image velocimetry) fields of cells around different division events. The PIV fields are rotated so that the long axis of the condensed chromosome is aligned along the *x*’-axis. The velocities are calculated at a time scale of 5 min. Scale bar 20 μm. The colorbar represent velocity divergence which scales linearly from −1.5×10^-3^ to 3×10^-3^ min^-1^. (h) 2D spatial pattern of ⟨*δd*(*δt*)⟩ at the division moment averaged over different division events. The patterns of ⟨*δd*(*δt*)⟩ are rotated so that the long axis of the condensed chromosome is aligned along the *x*’-axis. The inset shows the one without rotation. Scale bars 10 μm. (i) Spatiotemporal evolution pattern of ⟨*δd*(*δt*)⟩. The color bar scales linearly from 0.3 μm to 2 μm in (h) and from 0.3 μm to 3 μm in (i).

**Fig. 4.**
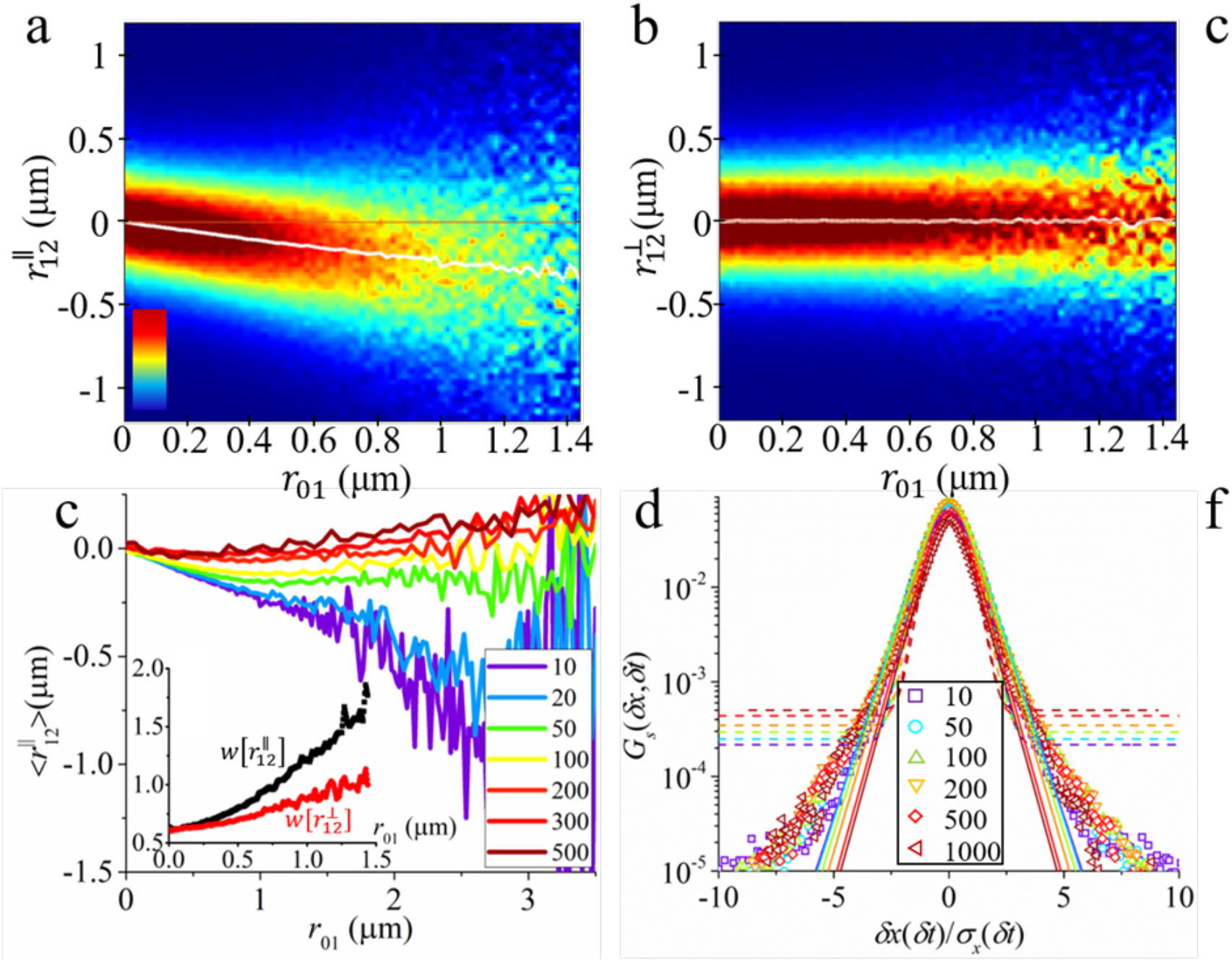
Creep motion of individual cells. (a) and (b) Conditional probabilities 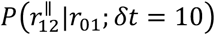 and 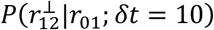(see text for definition). The red solid lines are guides for eyes showing 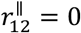 and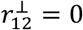. The white lines represent the mean values of 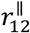 and 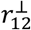 as a function of *r*_01_. The color bar represents the probability density which scales linearly from 0 (dark blue) to 0.05 μm^-1^ (dark red). (c) The mean values of 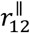 at different *δt* as a function of *r*_01_. Inset: the FWHM of 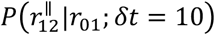 (black) and 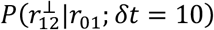 (red) shown in (a) and (b) as a function of *r*_01_. (d) Self-part of the Van Hove function for *δx* at different *δt*. The dashed and solid lines are fits with a Gaussian and a Gumbel law, respectively.

**Fig. 5.**
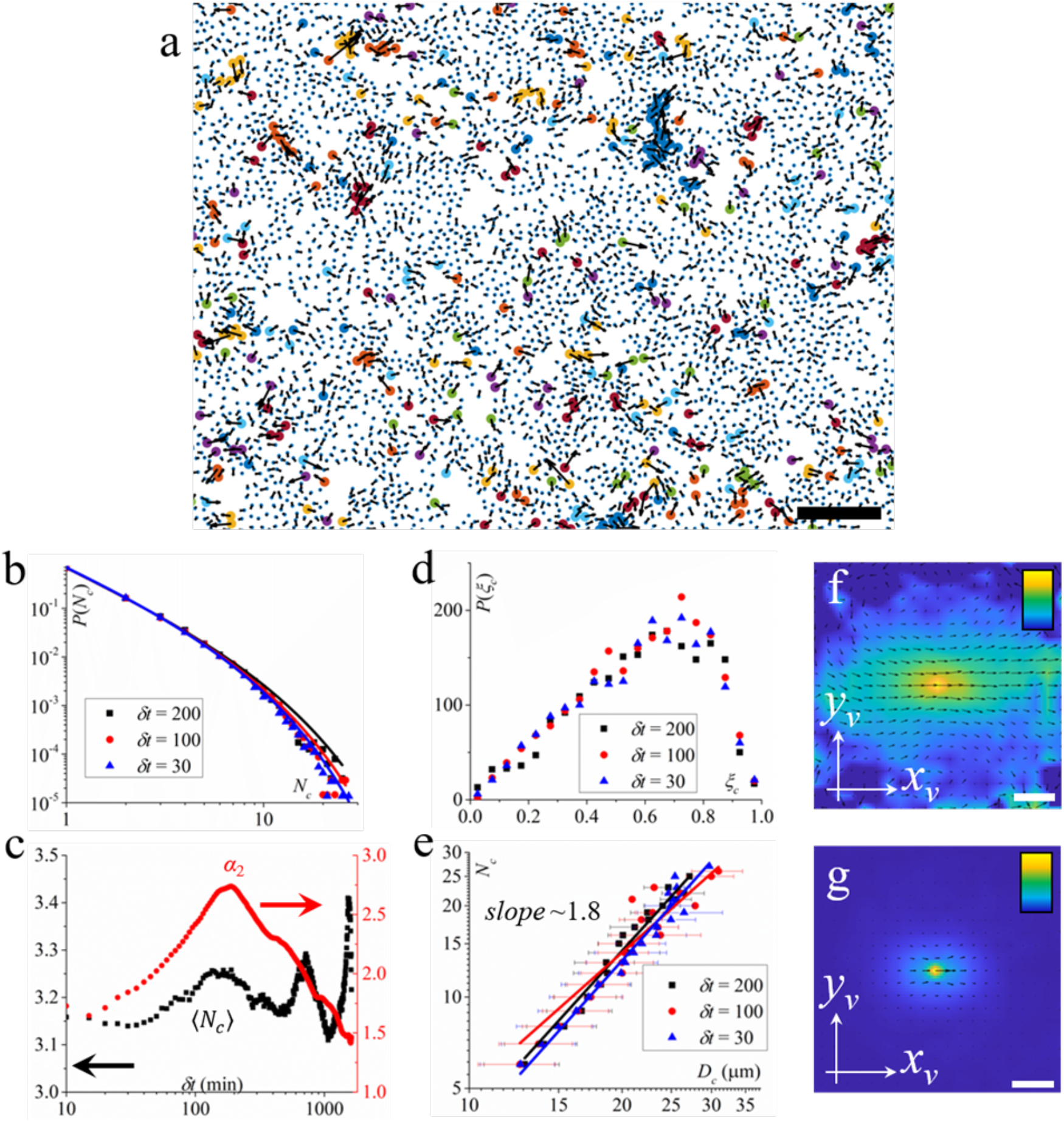
Dynamics and structure of fast cellular clusters. (a) A snapshot showing the dynamics of cells. The black arrows represent the displacements of cells at *t*+*δt* (*δt* =200 min), which are magnified by 3 times for a better view. The small blue dots represent the positions of cells at *t*. The large circles of different colors represent different cellular clusters that move quickly (top 10%). The scale bar is 100 μm. (b) PDF (probability density function) of cluster size, *N*_*c*_, for different time *δt*. (c) Mean size of cellular clusters <*N*_*c*_> (black) and non-Gaussian parameter *α*_2_ (red) as a function of time step *δt*. (d) PDF of anisotropy of cellular clusters, ξ_*c*_. (e) Double logarithmic plot of cluster size *N*_*c*_ as a function of cluster diameter *D*_*c*_. The solid black, red and blue lines are power law fits having exponents 1.84, 1.58 and 1.86, respectively. 2D velocity correlation functions of cells in fast clusters (f) and all cells (g) in a local coordinate system where the velocity of the reference cell is always aligned along the positive direction of the *x*_*v*_-axis. The color bars scale linearly from 0 to 1. Scale bars are 20 μm. The black arrows represent the mean velocity field.

**Fig. 6.**
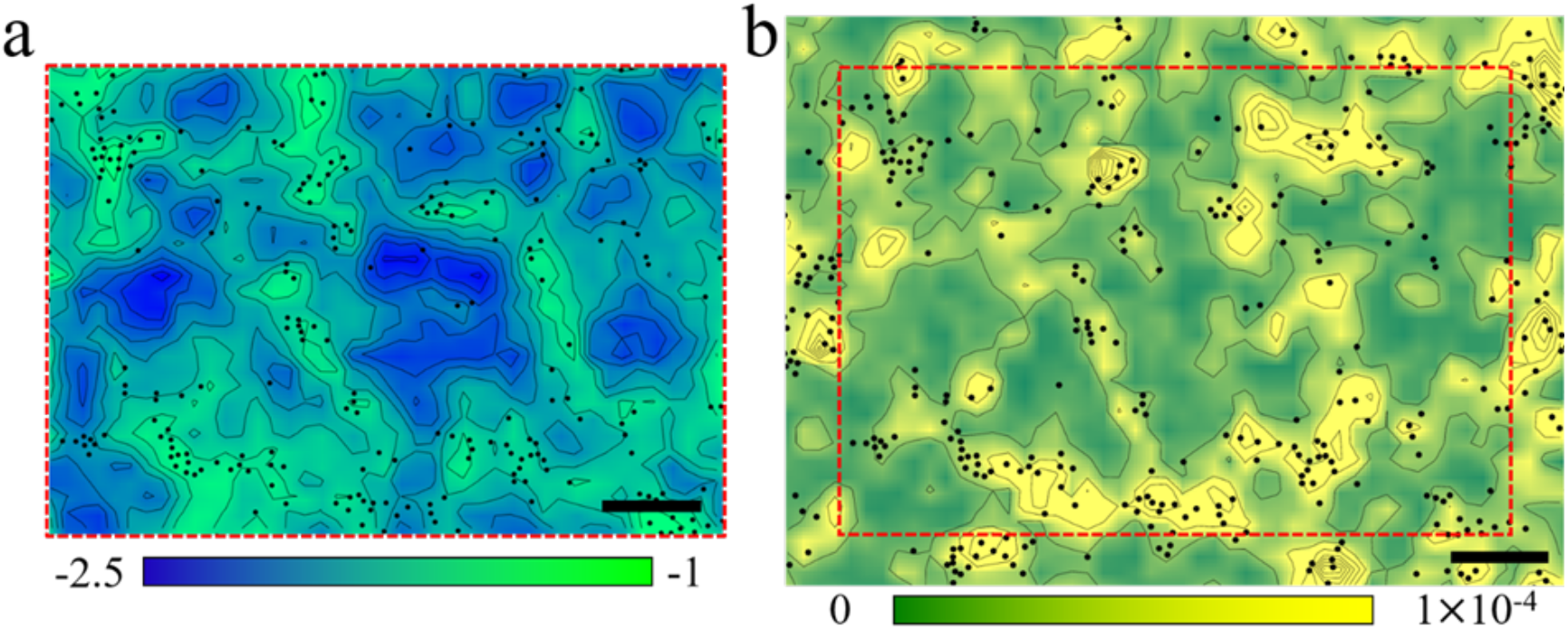
Spatial correlations between local properties and cell mobility. Contour maps of local structural entropy, *S*, (a) and mean magnitude of low-frequency normal modes, <*M*>, (b). The black dots represent the mobile cells with the 10% smallest values of self-overlap parameter *Q*_*i*_(*a*, *δt*). The color bars scale linearly from −2.5 to −1 for *S* and from 0 to 1×10^-4^ for <*M*>. Scale bars are 100 μm. The contour map of *S* is only calculated within the red dashed square region of the contour map of <*M*> to exclude the boundary effect.

The anisotropic shape of cell nuclei suggests that cellular motion is not isotropic and Fig. S3c demonstrates that this is indeed the case. The inset in Fig. 2b shows the diffusion coefficient *D* as a function of the direction with respect to the long axis of cell nuclei, (*D* is obtained in the diffusive regime, *δt* ∼ 200 to 400 min, of the TMSD according to the Einstein relation *δr*^2^(*δt*) = 4*Dδt*). The results show anisotropic diffusion, with a higher diffusion coefficient along the nuclear long axis. To further investigate the dynamics in the monolayer, we analyze the incoherent intermediate scattering function *F*_*s*_(*q*, *δt*) = ⟨exp(*iq*Δ*r*_*j*_)⟩ where Δ*r*_*j*_ = |**r**_*j*_(*δt*) − **r**_*j*_(0)| represents the displacement of cell *j* during the time interval Δ*t, q* is the wavenumber (*13*). We find, Fig. 2c, that this function decays in a simple manner, does not display the two-step relaxation that indicates the presence of cage effects (*13, 30*). By fitting *F*_*s*_(*q*, *δt*) with a stretched exponential function, 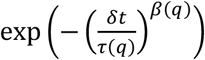, where *τ*(*q*) represents the relaxation time, we find that *τ*(*q*) exhibits a power-law dependence on the wavevector, *τ*(*q*) ∼*q*^−1.57^, Fig. 2c Inset, deviating from the *q*^-2^ scaling expected for Gaussian diffusion. This indicates that, while the system displays Fickian diffusion, i.e., the MSD at large times grows linearly, the underlying displacement distribution is not a Gaussian. Such “Fickian but non-Gaussian” dynamics has recently attracted significant attention across a variety of complex systems, including crowded intracellular environments, glassy materials, non-equilibrium systems as well as fluctuations in finance and politics (*31-33*). Our results suggest that such unexpected dynamics, possibly arising from dynamical heterogeneity or slow but intermittent active fluctuations, govern the relaxation behavior in jammed epithelial layers. At the same time, we find that at small *q*, the stretching exponent *β*(*q*) exceeds 1.0 (Fig. 2c inset), indicating a compressed exponential decay of *F*_*s*_(*q*, *δt*). Such compressed relaxations are often associated with internal stress relaxation processes, where collective rearrangements drive faster-than-diffusive dynamics (*34*). This is consistent with the presence of mechanical stresses in the jammed epithelial system (*4*), where force transmission between neighboring cells can induce coordinated relaxation events. Similar compressed behaviors have been broadly observed in soft glassy materials (*35-37*).

In contrast to the translational dynamics of the nuclei, the rotational dynamics is sub-diffusive over the whole-time scale of the experiment (Fig. S3b). This persistent memory of the orientation of the cell is likely one of the reasons why their translational dynamics is Fickian but non-Gaussian for surprisingly long times.

### Influence of cell divisions and extrusions

Previous theoretical models have suggested that cell divisions and apoptosis might fluidize jammed epithelial tissues (*26, 27*). To test whether the absence of cage effect is due to these mechanisms, we analyze the dynamics of cells surrounding division and extrusion events (Fig. 3 for divisions, S4 for extrusions). For this we quantified the time dependence of cell displacement, *d*, averaged over cells in a square region centered on a division [extrusion] event (Fig. 3a, b and S4a, b). The size of the square region is 100 × 100 μm^2^, which is large enough to have good statistics and also small enough to clearly see the spatiotemporal influence of cell divisions [extrusions]. We define as <*d*_div_> [<*d*_ext_>] as the mean cell displacement averaged over different division [extrusion] events (*Methods*) and comparing it to the one of random control regions of same size, <*d*_ran_>. Figures 3c and S4c demonstrate that <*d*_div_> [<*d*_ext_>] is close to <*d*_ran_> before cell division (*δt* < 0) but becomes larger than <*d*_ran_> immediately after division [extrusion] (*δt* > 0), indicating a local enhancement of cell motility around division [extrusion] events. A double-logarithmic plot of <*d*_div_> [<*d*_ext_>] and <*d*_ran_> reveals that both exhibit the same subdiffusive behavior (insets in Fig. 3e and S4e), suggesting that division [extrusion] does not change the type of cell motion in a significant manner.

This enhancement of cell mobility is also observed in the mean displacement step, ⟨*δd*(*δt* = 10)⟩ (*Methods*), which peaks near the division/extrusion time and decays rapidly within ∼30 min (Fig. 3d and S4d), confirming that the transient boost in mobility is short lived. Also the visualization of the averaged velocity fields supports this transient, collective motion (Fig. 3f, g and S4f, g). To assess its spatial extent, we examined how ⟨*δd*(*δt*)⟩ varies with distance from the division [extrusion] center. We find that the boost effect is confined to ∼20 μm (approximately 2 to 3 cell diameters), i.e., it is strongly localized in space (Fig. 3e, h and S4 e, h). Both the spatial and temporal limits of this boost effect triggered by cell divisions [extrusions] can be well visualized from the spatiotemporal evolution pattern of ⟨*δd*(*δt*)⟩ as shown in Fig. 3i and S4i.

From the above results one thus concludes that cell divisions and extrusions can indeed fluidize the jammed state of epithelial tissues. However, this effect is both spatially localized and short-lived. In addition, these events are relatively rare at high cell densities: Among ∼4,700 cells, we observe on average only one division every ∼10 minutes and one extrusion every ∼40 minutes. Given their low frequency and localized nature, we conclude that they are too sparse and too weak to account for the global absence of caging effects or to drive large-scale fluidization in jammed epithelia, which agrees with a theoretical model reported before (*22*).

### Creep motion of individual cells

To understand the identified fluidity of the jammed tissues, we consider the function 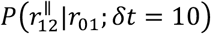 which is the conditional probability of the projection of the cell displacement in the time interval [*δt*, 2*δt*] (given by 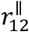) along the direction of the previous cell displacement in the time interval [0, *δt*] (given by *r*_01_). (In the same way one can define the conditional probability 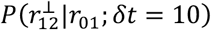 (*28, 38*).Figure 4a demonstrates that the mean value of 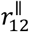 decreases linearly from 0 to negative values as *r*_01_ increases, indicating the presence of memory effects: cells that have moved in the direction **r**_01_ during the time interval [0, *δt*] are more likely to move in the opposite direction during the time interval [*δt*, 2*δt*], an effect that is reminiscent of elastic behavior. In addition, it is noted that the full width at half maximum (FWHM, *Methods*) of 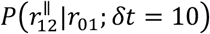 increases with *r*_01_ which indicates that larger (backward) displacements are more likely for cells which moved farther in the previous time interval (Fig. 4c inset). On the other hand, the mean value of 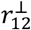 remains centered around 0 (Fig. 4b) and the FWHM of 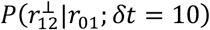 shows minimal dependence on *r*_01_ (Fig. 4c inset), suggesting there is no correlation between 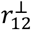 and **r**_01_. These results indicate that cells undergo a microscopic creep motion characterized by directionally anti-correlated steps, which may partially explain the observed sub-diffusion on short time scales. In addition, it is found that as *δt* increases, the slope of the mean value of 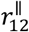 gradually approaches 0 (Fig. 4c). This demonstrates that the memory effect gradually decreases with *δt* and vanishes when *τ* ∼ 200 min. This time scale is consistent with the crossover of the TMSD from sub-diffusive to diffusive regime (Fig. 2a). We also note that the mean value of 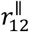 becomes even positive at large *r*_01_ when *δt* is large. This may be attributed to a local collective motion of cells which allows them to move quickly over a distance of several cell diameters. Understanding the details of this accelerated dynamics is likely important for cell migration on the mesoscopic scale, work that should be done in future studies.

More details about the microscopic relaxation dynamics of the jammed tissue can be obtained from the probability density function (PDF) of displacements, also known as the self-part of the Van Hove function, *G*_*s*_(*δx*, *δt*). Fig. 4d shows the PDF of cell displacements along the *x*-axis, *2x*, of different time steps, *δt*. The displacements are rescaled by the root mean square displacements, 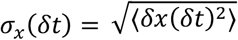.We observed that the rescaled curves collapse onto a single, conserved profile which cannot be well fitted with normal distribution, but can at small and intermediate displacements be well described by a Gumbel law 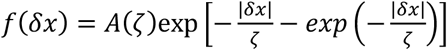,where *ζ* is a length scale and *A*(*ζ*) is a normalization constant (*39*). At larger displacements, *G*_*s*_(*δx*, *δt*) displays a significant excess tail with respect to the Gumbel law, indicating an enhanced number of relaxation events on larger length scales (*40*). Interestingly, we find that *G*_*s*_(*δx*, *δt*) does not approach the Gaussian distribution even at large time scales, which further confirms the presence of the Fickian-non-Gaussian behavior. Together with the memory effects, our results highlight the intermittent, anisotropic nature of motion in jammed epithelial tissues, where cells undergo local creep interspersed with rare, larger rearrangements. Such creep motion at the single-cell scale is qualitatively distinct from the dynamics of thermal glass-forming systems, and instead more closely resembles the behavior of complex fluids.

### Collective motion of cellular clusters

In addition to the creep motion of individual cells at microscopic scales, we observe that the dynamics of the jammed tissue is strongly heterogeneous in that at a given time different regions relax with different rates (Fig. S5). Such a phenomenon is often called “dynamic heterogeneity” and has been widely observed in thermal glassy systems (*30, 41, 42*), colloidal suspensions (*43, 44*) and granular materials (*45, 46*). To understand the formation of this dynamic heterogeneity and its influence on the fluidity of the jammed tissues, we investigate the spatial organization of fast-moving cells. To do this, we identified the top 10% most mobile cells based on their displacement magnitude at different time difference *δt*. Using a nearest neighbor criterion with a cutoff distance given by the location of the first minimum in radial distribution function, see Fig. S6 and *Methods*, we can group these fast cells in distinct clusters (Fig. 5a). Figure. 5b shows the PDF of cluster sizes (the number of fast cells in a cluster), *N*_*c*_, at three different *δt. P*_*c*_(*N*_*c*_) first decays as a power law and then exhibits an exponential tail which can be well described by 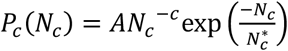, where 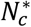 represents the cutoff size which characterizes the transition from power law to exponential behavior (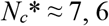, and 5 for *δt* = *200, 100 and 30* min, respectively.), *c* ≈ 1.8 is a fitting parameter, which is slightly smaller than the value of random percolation (*c* ≈ 2.1) (*47*) and thus, indicating the cooperative motion of cells within clusters. The pre-factor *A* is determined from the normalization condition 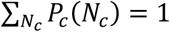.Similar cluster size distribution described here has also been reported in fish schools (*48*), buffalo herds (*49*), bacteria swarms (*50*) and many other animal species (*51*). Fig. 5c shows the dependence of the mean cluster size (number of fast cells), <*N*_*c*_>, on time, *δt*, and the temporal dependence of the non-Gaussian parameter, *α*_2_ (*Methods*), respectively. We find that <*N*_*c*_> gradually increases with *δt*, forming a peak at *δt*∼*200* min, and then decreases with *δt*. The strong oscillations at the late time stage is likely due to statistical errors caused by a few large persistent clusters. On the other hand, the non-Gaussian parameter, *α*_2_, which characterizes the deviation of cell displacements from a Gaussian distribution, shows a similar temporal dependence as <*N*_*c*_> and forms a peak at *δt*∼200 min, corresponding to the end of the sub-diffusive regime in the TMSD (Fig. 2a). *α*_2_ gets large when the cell displacement distribution has fat tails, i.e., when there is a pronounced subpopulation of “anomalously fast” cells at *δt*. Such a correlation between <*N*_*c*_> and *α*_2_ indicates that the motion of fast cells is cooperative. At large times, *α*_2_ decays to zero because the system approaches Gaussian dynamics, but that limit seems to be reached only on time scales of 10^4^ minutes, i.e., a time that is surprisingly long.

To characterize the geometry of fast-moving cell clusters, we computed their inertia tensor which quantifies how cell positions are spatially distributed within each cluster (see *Methods*). It allows to compute each cluster’s anisotropy, ξ_*c*_, and characteristic diameter, *D*_*c*_. ξ_*c*_ = 1 for a linear structure and ξ_*c*_ = 0 for a disk-like structure (*52*). These metrics allow us to quantify whether fast-moving clusters are isotropic blobs or anisotropic, string-like structures — a key question in understanding how collective rearrangements propagate in jammed tissues. It should be noted that here only cellular clusters with *N*_*c*_ > 5 are counted. Fig. 5d shows the PDF of ξ_*c*_ for 3 different *δt*. They form a peak at ξ_*c*_∼0.7, indicating an elongated shape of the clusters. In addition, the radial distribution function suggests that these clusters are elongated along their mean velocity direction (Fig. S7). Furthermore, the relationship between *N*_*c*_ and *D*_*c*_ reveals that fast clusters exhibit self-similar, fractal-like behavior, 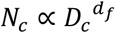,where *d*_*f*_ ≈ 1.8 denotes the Hausdorff dimension (Fig. 5e). This value is close to the *d*_*f*_of diffusion-limited aggregates (DLA, *d*_*f*_ ≈ 1.7) (*53*) and the *d*_*f*_ of critical percolation clusters (*d*_*f*_ ≈ 1.89) (*54*), indicating these cellular clusters have a ramified, porous structure which is more compact than DLA but a bit looser than critical percolation.

Finally, to understand the motion of cells within clusters, we measure the velocity correlation function, *C*_*v*_(*δr*) = ⟨**v**_*i*_(0)**v**_*j*_(*δr*)⟩, where **v** is cell velocity and *δr* represents the distance between cell pairs, of cells within large fast clusters (*N*_*c*_ > 5) (Fig. 5f) and compare it with the one of the whole system (Fig. 5g). Here, both correlation functions are calculated in a local coordinate system where the velocity of the reference cell is always aligned along the positive direction of the *x*_*v*_-axis. As we can see, *C*_*v*_ of the fast clusters is much more long-ranged compared to the one of the whole system, indicating cells move coherently within clusters consistent with our conclusions from the correlation between <*N*_*c*_> and *α*_2_ (Fig. 5c). In addition, *C*_*v*_ exhibits a stronger long-range correlation along the mean cluster velocity, which is consistent with the anisotropic structure of the clusters (Fig.S7).

Taken together, these results indicate that fast-moving cells in jammed tissues are not isolated, but self-organize into elongated, collective structures. Such clusters may act as conduits for stress relaxation or as transient rearrangement pathways, enabling tissues to locally remodel without full unjamming. This collective behavior could be particularly relevant during processes like wound healing or morphogenesis, where localized reorganization occurs within globally arrested environments.

### Microscopic origin of dynamical heterogeneity

Having established that fast-moving cells form collective, anisotropic clusters, we now investigate the origin of this dynamical heterogeneity, i.e., why do certain cells undergo large displacements and rearrangements while others remain basically static? Is this variability random, or does it reflect structurally distinct “soft spots”, akin to dislocations in crystalline solids (*55*), that are predisposed to rearrangement? While we have shown that cell divisions and extrusions can locally accelerate cellular motion, their effects are spatially localized and temporally transient. Thus, these active events alone cannot account for the widespread heterogeneity observed in the tissue.

To address this question, we quantify the degree of local rearrangement using the self-overlap parameter *Q*_*i*_(*a*, *δt*), which measures the similarity of two configurations separated by a time *δt*, on a length scale set by *a* (see *Methods* for the definition of *Q*_*i*_(*a*, *δt*)). Unless otherwise noted, we use *a*=1.2 μm (which is about 10% the size of a single cell) as the reference length scale and *δt*=200 min as the time step to capture meaningful single-cell rearrangements at large time scales. (The temporal dependence of *Q*_*i*_(*a*, *δt*) of different values of *a* is presented in Fig. S8.) We then calculate the correlation of *Q*_*i*_(*a*, *δt*) with various structural descriptors: local cell density, *ρ*_*i*_, local nematic orientational order of nuclei, 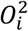, and local hexatic bond orientational order of nuclei, 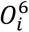 (*Methods*). Spatial maps (*Methods*) and Pearson correlation coefficients reveal negligible correlations between rearrangements and these standard order parameters (Fig. S9), demonstrating that local density and geometric order alone do not predict dynamic activity.

In contrast, we find that local structural entropy, *S*_*i*_ (*Methods*), which captures the structural disorder in a cell’s local environment (*56*), shows a moderate negative correlation with *Q*_*i*_(*a*, *δt*) (the Pearson correlation coefficient is *P* ≈ −0.43). Cells that are persistently in high-entropy regions are more likely to undergo rearrangements (Fig. 6a), indicating that local structural disorder promotes fluidity, similar to what has been reported in thermal glassy systems (*56, 57*). Building on this, we examined whether dynamic heterogeneity is also linked to low-frequency vibrational modes, as reported in glassy systems (*58, 59*). We computed the displacement covariance matrix from early-time dynamics and extracted the spatial distribution of the lowest-frequency modes (*60, 61*). It should be noted that the measurements of the soft modes and *Q*_*i*_(*a*, *δt*) are separated in time (*Methods* for details). Remarkably, cells located in regions of large vibrational amplitude are more likely to undergo rearrangements later (Fig. 6b) as demonstrated by the negative Pearson correlation coefficient (*P* ≈ −0.36), between the mean magnitude of low-frequency normal modes, <*M*>, and *Q*_*i*_(*a*, *δt*). This demonstrates a predictive link between soft vibrational modes and subsequent cellular dynamics, suggesting mechanical susceptibility — region’s propensity to undergo rearrangement, remains encoded in the tissue’s structure.

Together, these findings indicate that dynamic heterogeneity in jammed epithelia arises not from local density or geometric order, but from subtle variations in structural disorder and local soft vibrational modes. Considering the strong stress fluctuations in epithelial tissues reported before (*4, 62*) and the stress driven relaxation behavior discussed above (Fig. 2c), the correlation observed here may be attributed to the large stress accumulation in regions of high structural disorder and soft vibrational modes — an intriguing hypothesis that warrants future investigation.

## Discussion

In this work, we provide a comprehensive experimental characterization of the single-cell dynamics in jammed epithelial tissues. Our central finding is that high-density epithelial tissues are not static/rigid as thermal glassy systems. Instead, individual cells undergo sub-diffusive creep motion that gradually transitions into Fickian yet non-Gaussian diffusion. This behavior suggests that even in a jammed state, epithelial tissues remain fluid-like at the microscopic scale, continuously yielding and rearranging. Rather than being mechanically frozen, epithelia flow as complex fluids that cells explore their local environments through persistent, low-amplitude motion. These findings fundamentally revise the notion that high-density epithelia are rigid or inert, but instead align with recent simulation studies on active glass showing that persistent active forcing can fluidize dense systems without requiring cage escape or glassy relaxation dynamics found in thermal glass-formers (*63, 64*).

We further show that cell divisions and extrusions serve as local, transient sources of mechanical activity. Although they briefly accelerate the motion of neighboring cells, their influence is spatially confined and short-lived. This indicates that tissue-wide fluidity is not solely driven by active biological events but is instead an emergent feature of the tissue’s structural and mechanical organization.

Beyond the absence of caging, the system exhibits unexpected features in its displacement statistics. While cell motion is diffusive at long times, the displacement distributions remain strongly non-Gaussian, a behavior known as “Fickian but non-Gaussian” diffusion. Such a dynamics is characteristic of systems with heterogeneous relaxation pathways, such as crowded intracellular environments or active soft materials (*31-33*). Moreover, we observe compressed exponential decays in the incoherent intermediate scattering function at low wavevectors, a signature of internal stress relaxation via collective rearrangements. These features underscore the role of mechanical heterogeneity and stress propagation in shaping epithelial dynamics, highlighting parallels with soft glassy systems (*35-37*).

Our identification of fast-moving, anisotropic clusters of cells provides further evidence for dynamic heterogeneity. These clusters are spatially coherent and elongated along the direction of motion, resembling string-like cooperative rearrangements seen in granular and colloidal systems (*45*). Importantly, we find that the occurrence of such clusters is correlated with local structural entropy and low-frequency vibrational modes — features long associated with “soft spots” in passive disordered materials (*56, 58*). This suggests that epithelial tissues possess regions predisposed to rearrangement, even in the absence of external perturbations. From a biological perspective, our findings reveal that jammed tissues retain an intrinsic capacity for reorganization, i.e., a hidden fluidity that may be essential for processes such as tissue morphogenesis, wound repair, and tumor progression. For instance, during development, epithelial sheets often undergo large-scale deformation without overt unjamming. The capacity for localized rearrangement may allow tissues to respond to morphogen gradients or mechanical cues while preserving global cohesion. Similarly, in early carcinogenesis, subpopulations of cells may leverage this residual plasticity to initiate invasive behavior before full-scale epithelial-to-mesenchymal transition (EMT) occurs.

Altogether, our study bridges concepts from soft matter physics and epithelial biology to uncover how dense tissues sustain microscopic motion and mechanical responsiveness. By quantifying the interplay between structural disorder, internal fluctuations, and dynamic heterogeneity, we propose a new framework for understanding the emergent mechanics of jammed epithelia. Although our findings are based on MDCK cells, the conserved structural organization of epithelial tissues suggests that similar behaviors may be present in other systems — a hypothesis warranting future investigation. These results open the door to studying tissue remodeling and soft matter-like fluidity in both physiological and pathological contexts.

## Methods

### Cell lines and culturing conditions

Madin-Darby Canine Kidney (MDCK) cells stably expressing histone1-GFP were cultured in glass-bottom petri dishes (FluoroDish, catalog no. FD35-100) for at least two weeks to ensure the epithelial monolayer reached a deeply jammed state. Cells were maintained in Dulbecco’s Modified Eagle Medium (DMEM, containing glucose and pyruvate; Life Technologies) supplemented with 10% fetal bovine serum (FBS; Life Technologies) and 1% penicillin-streptomycin (Life Technologies), with medium changes every two days. To prepare the substrate, the glass surface of each petri dish was first plasma-cleaned for 3 minutes, then coated with a layer of polymerized Matrigel. To initiate polymerization, EDC (N-(3-Dimethylaminopropyl)-N′-ethylcarbodiimide hydrochloride, E1769 Sigma-Aldrich) and NHS (N-Hydroxysuccinimide, #130672 Sigma-Aldrich) were dissolved in cold calcium- and magnesium-free PBS and mixed with Matrigel to achieve final concentrations of 40 mM EDC and 10 mM NHS. About 50 µL of this mixed solution was smeared on the cleaned glass-bottom petri dishes and incubated at 37 °C for 2 hours to allow complete polymerization. After polymerization, the coated dishes were rinsed three times with 1x PBS and incubated in 1x PBS at 37 °C before cell seeding. To ensure the reproductivity of our results, the measurements of MSD, incoherent intermediate scattering function, conditional probability distribution and Van Hove function, were repeated in 2 independent samples, which both show similar behaviors regarding their static and dynamic properties.

### Live cell imaging and data analysis

Samples were observed through a 10 × objective on a BioStation IM-Q (Nikon, Tokyo, Japan) at 37 ^°^C and 5% CO_2_ with humidification. Images were taken every 5 min for about 27 hours. More than 3800 cells were tracked over a field of view of 828 μm × 621μm. It should be noted that there were overall about 4700 cells in the field of view, but around 900 of them whose trajectories were lost during the tracking over a time period of 27 hours. So, we only analyzed the rest 3800 cells whose trajectories went through all the frames. The movies were then analyzed through Cellpose2.0 ^55^, ImageJ and MATLAB. Basically, nuclei were first segmented by Cellpose, which were then tracked using the Trackmate plugin of ImageJ. The tracking results (including the spatial coordinates of the center of mass of cell nuclei and the long-axis orientation of cell nuclei) were then analyzed through MATLAB.

To characterize the tracking error produced during the experiment, we compare the cell displacements of the jammed system with a fixed sample where cells are dead and thus do not move as shown in Fig. S1. It is found that the tracking error is basically independent of time and is around 0.1 μm, which is less than the half size of the cell displacements at the smallest *δt*. At the same time, cell nuclei can change their shapes over time. To characterize its influence, we calculate the mean values of the changes of the long- and short-axes of cell nuclei as a function of time, 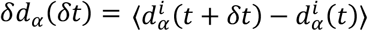, where 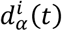 represents the length of the long (*α* = *l*) or short (*α* = *s*) axis of nucleus *i* at time *t*, ⟨… ⟩ represents averaging over different cell nuclei at different times. It is found that both of them are much smaller compared to the cell displacements and become negligible at large *δt*. So, we conclude that the tracking error of our experiments is around the half length of cell displacements at the smallest *δt* and becomes negligible as *δt* increases.

The TMSD is calculated as ⟨*δ***r**^2^(*δt*)⟩ = ⟨(**r**_*i*_(*t* + *δt*) − **r**_*i*_(*t*))^2^⟩, where **r**_*i*_(*t*) is the position of cell *i* at time *t*. The cage-related TMSD is defined as, 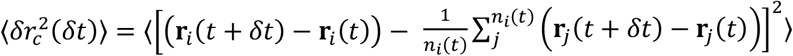, where *n*_*i*_(*t*) is the number of nearest neighbors of cell *i* at time *t*, the nearest neighbors are defined as cells whose distances between themselves and the reference cells is smaller than the distance of the first peak of the radial distribution function of the system.

The mean displacements of cells over different division [extrusion] events is defined as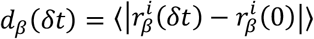, where 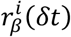 represents the spatial coordinate of cell *i* in the division (*β* = *div*) / extrusion (*β* = *ext*) region at time *δt* away from division [extrusion] moment, ⟨… ⟩ represents average over different local cells around division [extrusion] events, *δt* represents the time delay before and after the division [extrusion] moment. The mean displacement step is defined as 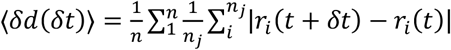, where **r**_*i*_(*t*) represents the coordinate of cell *i* around division [extrusion] event *j* at time *t, δt* =10 min represents time interval, *t* represents the time moment away from the division moment, *n*_*j*_ and *n* represent the number of cells around division event *j* and the number of division events, respectively.

To calculate the FWHM of 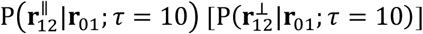,we first calculate the second moment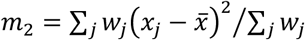, where 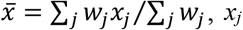represents the values of 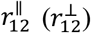 at specific *r*_01_, and *w*_*j*_ represents the corresponding probability weight of *x*_*j*_. The FWHM is then defined as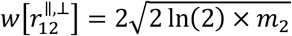.

To define the clusters of fast cells, we first calculate the cell displacements at each time as *d*_*i*_(*t*, *δt*) = |*r*_*i*_(*t* + *δt*) − *r*_*i*_(*t*)| and define the top 10% of cells having the largest *d*_*i*_(*t*, *δt*) as fast cells. We then define a threshold distance *r*\*=15 μm, which is the distance of the first valley of the radial distribution function of the epithelial system (Fig. S6). Two fast cells are considered to be neighbors if their intercellular distance is smaller than *r*\* and a fast cell belongs to a fast cellular cluster if it is the neighbor of any other fast cells belonging to the cluster.

The inertia tensor is defined according to refs. (*52, 65*), 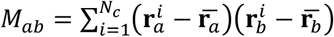, where *a, b* represents 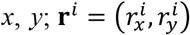 are the spatial coordinate of the *i*th cell; and 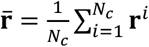 is the center of mass of the cluster. *M*_*ab*_ allows to define a characteristic diameter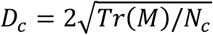,and anisotropy of the structure, 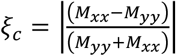.

The non-Gaussian parameter which quantifies the deviations from a Gaussian distribution is defined as 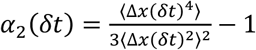, where Δ*x*(*δt*) is the cell displacement along the *x*-axis during the time interval *δt*.

The self-overlap parameter is defined as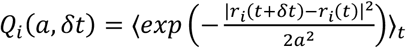, where *i* represents the specific cell, *a* is the reference length scale. *Q*_*i*_(*a*, *δt*) decays from 1 to 0 as cells gradually move away from their initial reference positions. It should be noted that here *Q*_*i*_(*a*, *δt*) is averaged over time from *t* = 105 min to *t* = 1425 min to reduce the noise. The reason that we can do this time average is because the cells are not moving that much within this time span, i.e., they do not change their nature (say from fast to slow) as shown in Fig. 2a. So, the time average improves the statistics but does not wipe out the interesting information.

The local nematic order parameter (*66*) is defined as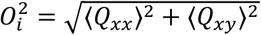, where *Q*_*xx*_ = *cos* (2*θ*) and *Q*_*xy*_ =*sin* (2*θ*) are components of nematic tensor *Q, θ* is the angle between the long axis of cell nuclei and a fixed axis, ⟨… ⟩ represents the average of the tensor *Q* over the reference cell *i* and its Voronoi neighbors.

The hexatic bond orientational order parameter is defined according to ref. (*67*), 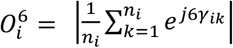, where *n*_*i*_ is the number of Voronoi neighbors of cell *j* and *γ*_*jk*_ represents the angle between a reference axis and the bond vector connecting cell *i* and its neighbor *k*.

The local structural entropy is defined according to ref. (*56*), 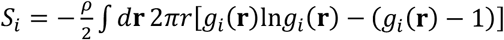, where *g*_*i*_(**r**) is the radial distribution function of cell *i*. Similar to *Q*_*i*_(*a*, *δt*), the contour maps of 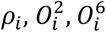 and *S*_*i*_ are averaged over the time period from *t* = 105 min to *t* = 1625 min to improve statistics.

The calculation of the vibrational modes can be found in refs. (*60, 68*). In brief, to find the low-frequency normal modes, we construct the covariance matrix, *D*_*kl*_ = ⟨*u*_*k*_*u*_*l*_⟩_*t*_, where *u*_*k*_ = *x*_*k*_ − ⟨*x*_*k*_⟩_*t*_, *y*_*k*_ − ⟨*y*_*k*_⟩_*t*_, *k, l* = 1,2, …,2*N* (*N* cells) run over all cells and their Cartesian components in two dimensions, based on the cell displacements in the first 100 minutes over which there are few cell rearrangements. We then obtain the eigenvalues *λ*_*k*_ and their corresponding eigenmodes **M**_*k*_. We average the magnitude of the 10 eigenmodes which have the 10 largest eigenvalues (lowest frequencies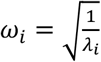) to get the contour map of mean magnitude of low-frequency normal modes, <*M*_*k*_>, and compare it with *Q*_*i*_(*a*, *δt*). It should be noted that here the self-overlap parameter, *Q*_*i*_(*a*, *δt*), is calculated based on the time window after the first 100 min, i.e. *T*_2_ = [105,1625] min, so there is no overlap between the calculations of **D** and *Q*_*i*_(*a*, *δt*).

To plot the colormap of 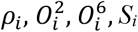 and <*M*>, we applied a coarse-graining method. Basically, we first divide the whole field of view of our data into small windows (40×40 μm^2^), with an overlap of 20 μm between the windows. Then, in each small window, we average the values of the specific parameter of the cells within that window. Finally, we interpolate those mean values and get the colormaps.

## Author Contributions

Y.S. conceived and carried out the experimental investigations, analyzed the results and wrote the first draft of the manuscript. W.X. contributed to the preparation of Matrigel substrates. All authors contributed to discussions and wrote the manuscript. W.K. and B.L. supervised the project.

## Acknowledgements

We thank the members of the “Cell adhesion and Mechanics” team for helpful discussions. This work was supported by the European Research Council (Grant No. Adv-101019835 “DeadorAlive” to BL), the Alexander von Humboldt Foundation (Alexander von Humboldt Professorship to BL), the Agence Nationale de la Recherche (“STRATEPI” DFG-ANR-22-CE92-0048 and 0048 and “VISCOMAG2” ANR-24-CE42-6142 to RMM), Institut National du Cancer (INCa_16712 and INCa_18429 to BL and RMM) and the “Initiatives d’excellence” (Idex ANR-11-IDEX-0005-02) transverse project BioMechanOE (TP5) (to W.X.) and Human Frontier Science Program (grant number LT0007/2023-C) (to YS). We acknowledge the ImagoSeine core facility of the IJM, a member of IBiSA and France-478 BioImaging (ANR-10-INBS-04) infrastructures.

## Data availability

The data that support the findings of the study are available from the corresponding authors upon reasonable request.

## Supplementary Figures

**Fig. S1.**
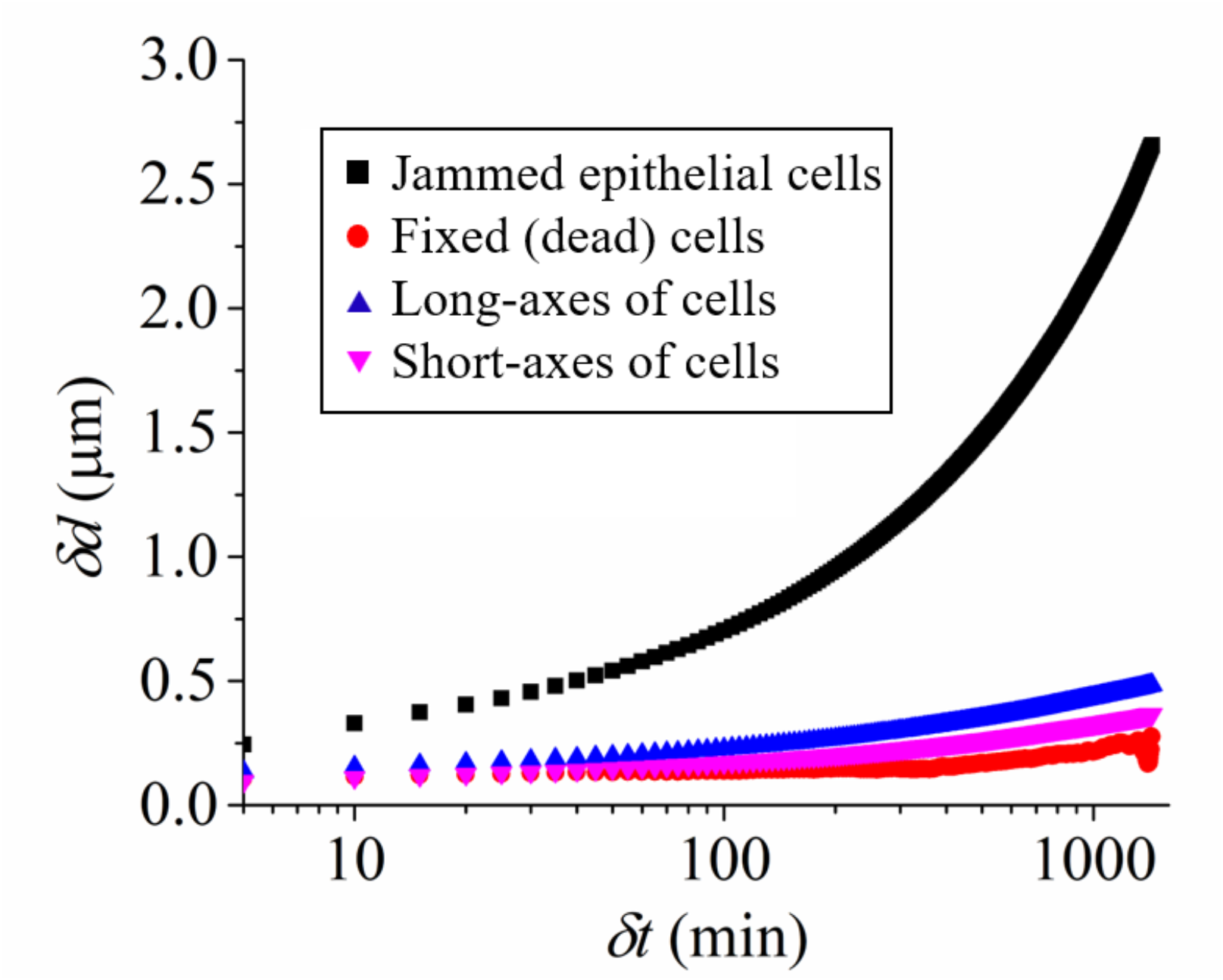
Cell displacement as a function of time delay. The black squares represent the displacements of the jammed epithelial sample in the manuscript. The red circles represent the displacements of fixed (dead) cells. Blue and pink triangles represent the changes of the lengths of the long and short axes of cell nuclei as a function of time delay. One recognizes that the displacement of the jammed cells is significantly larger than the one of the potential error sources.

**Fig. S2.**
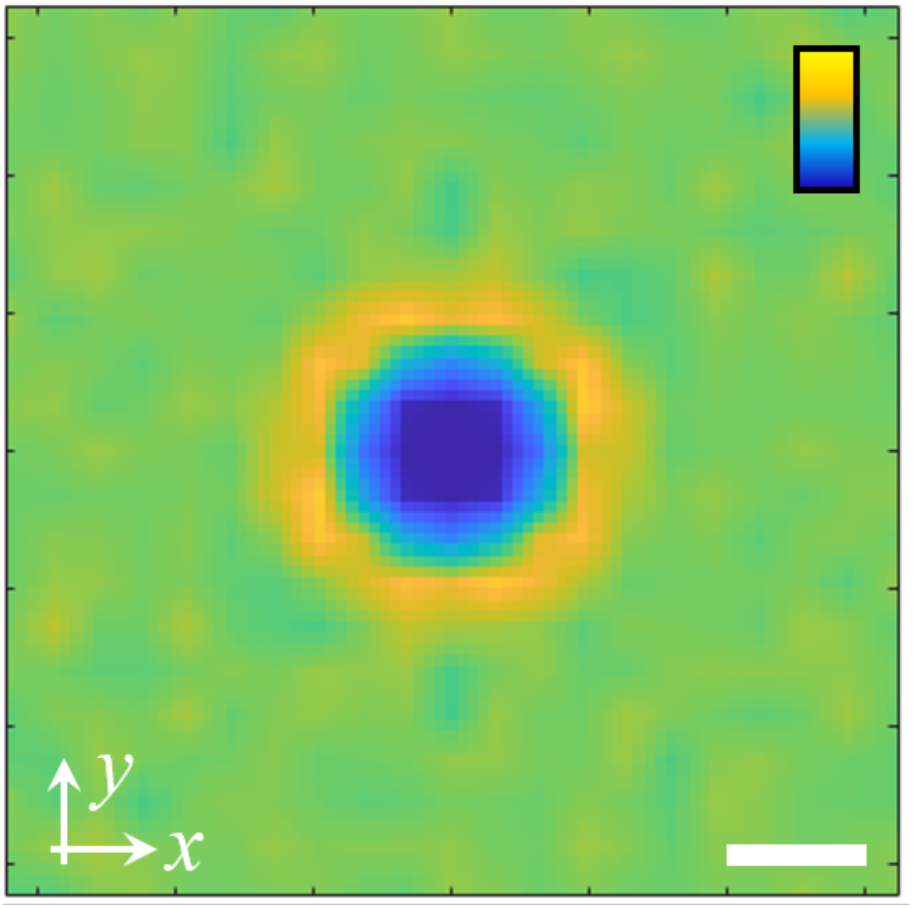
Pair correlation function of nuclei in the non-aligned lab coordinate frame. The scale bar is 10 μm. The color bar scales linearly from 0 (dark blue) to 1.5 (light yellow).

**Fig. S3.**
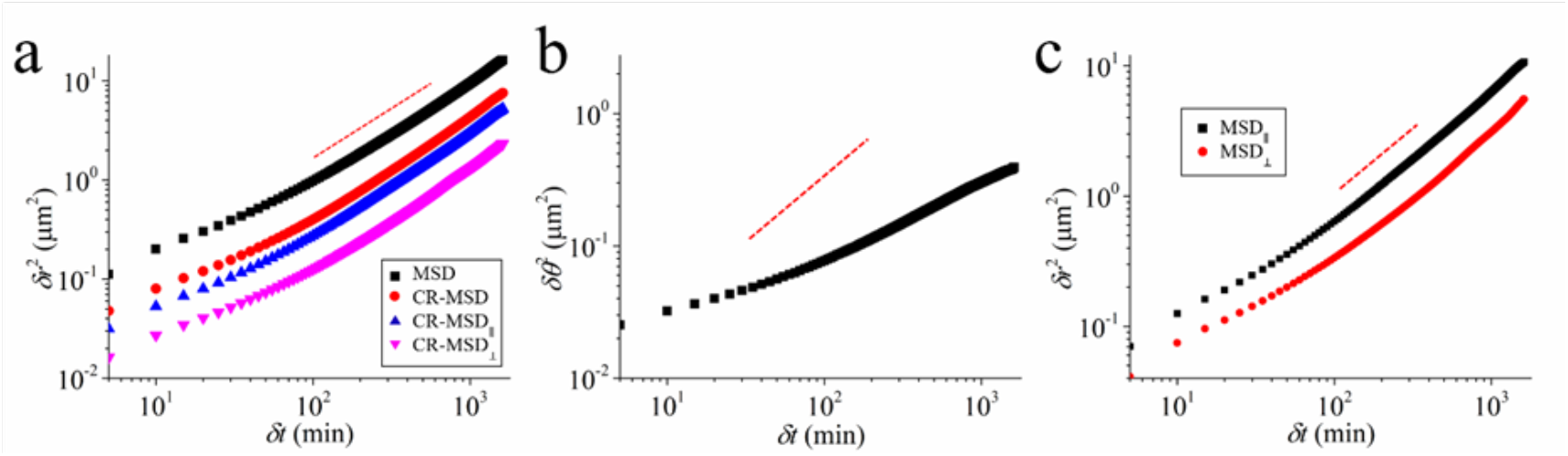
(a) Cage related TMSD of cells. The black squares represent the TMSD shown in Fig. 2(a) of the main text. The red circles represent the corresponding cage related TMSD. The blue and pink triangles represent the components of the cage related TMSD along and perpendicular to the long axes of cell nuclei, respectively. (b) Rotational mean squared displacement of cells as a function of time delay. (c) The TMSD of cells along (black) and perpendicular (red) to the long axes of cell nuclei. The red dashed lines in (a), (b) and (c) represent a power-law function with an exponent of 1.

**Fig. S4.**
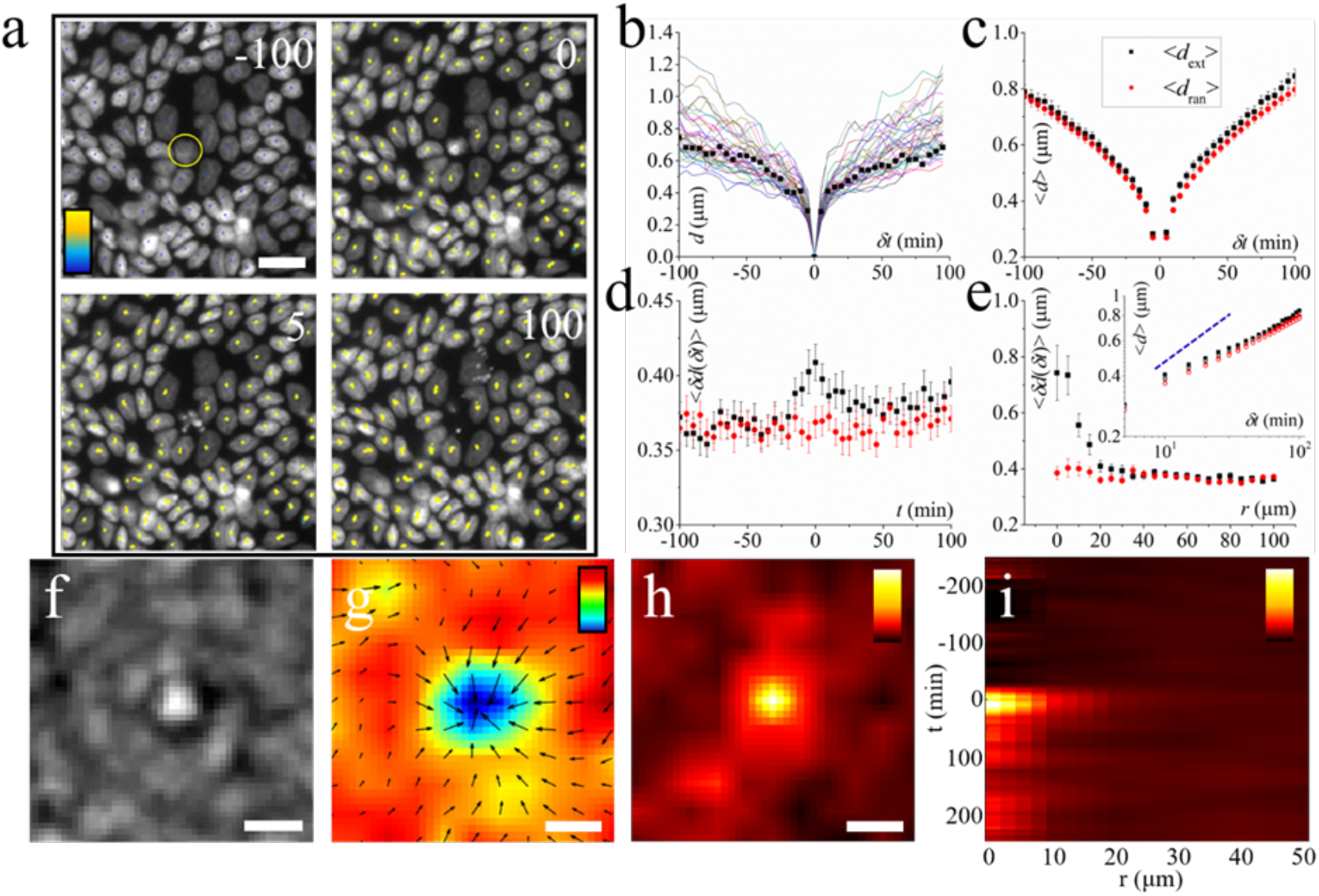
Dynamics induced by single cell extrusions. (a) Snapshots showing the extrusion of a cell at different moments. The dots represent cell trajectories. The color bar represents time lapse which scales linearly from 0 min (dark blue) to 200 min (light yellow). The extruding cell is marked with a yellow circle. The scale bar is 20 μm. (b) Cell displacement, *d*, averaged over local cells around an extruding cell as a function of time. *δt* represents time delay before and after extrusion. Solid lines of different colors represent cell displacements of different extrusion events. The black squares represent the cell displacement of the extrusion event in (a). (c) Mean cell displacement, <*d*>, averaged over different extrusion events (black squares, <*d*_ext_>) as a function of time. The red dots represent the mean cell displacement averaged over different random control regions, <*d*_ran_>. (d) Mean displacement step, ⟨*δd*(*δt*)⟩, of local cells averaged over different extrusion events (black squares) as a function of time before and after extrusion moment. The red dots represent ⟨*δd*(*δt*)⟩ of local cells in random square regions. (e) ⟨*δd*(*δt*)⟩ at the extrusion moment as a function of distance away from the extrusion event. The black (red) dots represent ⟨*δd*(*δt*)⟩ of local cells in extrusion (random) square regions. The inset shows the loglog plot of *d*_*ext*_(*δt*) (black) and *d*_*ran*_(*δt*) (red) in (c) as a function of time. The solid (hollow) symbols represent the mean cell displacements after (before) extrusion. The blue dashed line indicates a power law function with an exponent of 1/2. (f) Microscopic image obtained by averaging the brightfield images of different extrusion events. Scale bar 10 μm (g) Mean velocity field obtained by averaging the PIV (particle image velocimetry) fields of cells around different extrusion events. The velocities are calculated at a time scale of 5 min. Scale bar 20 μm. The colorbar represent velocity divergence which scales linearly from −1×10^-3^ to 3×10^-3^ min^-1^. (h) 2D spatial pattern of ⟨*δd*(*δt*)⟩ at the extrusion moment averaged over different extrusion events. Scale bars 20 μm. (i) Spatiotemporal evolution pattern of ⟨*δd*(*δt*)⟩. The color bar scales linearly from 0.3 μm to 0.8 μm in (h) and from 0.3 μm to 1 μm in (i).

**Fig. S5.**
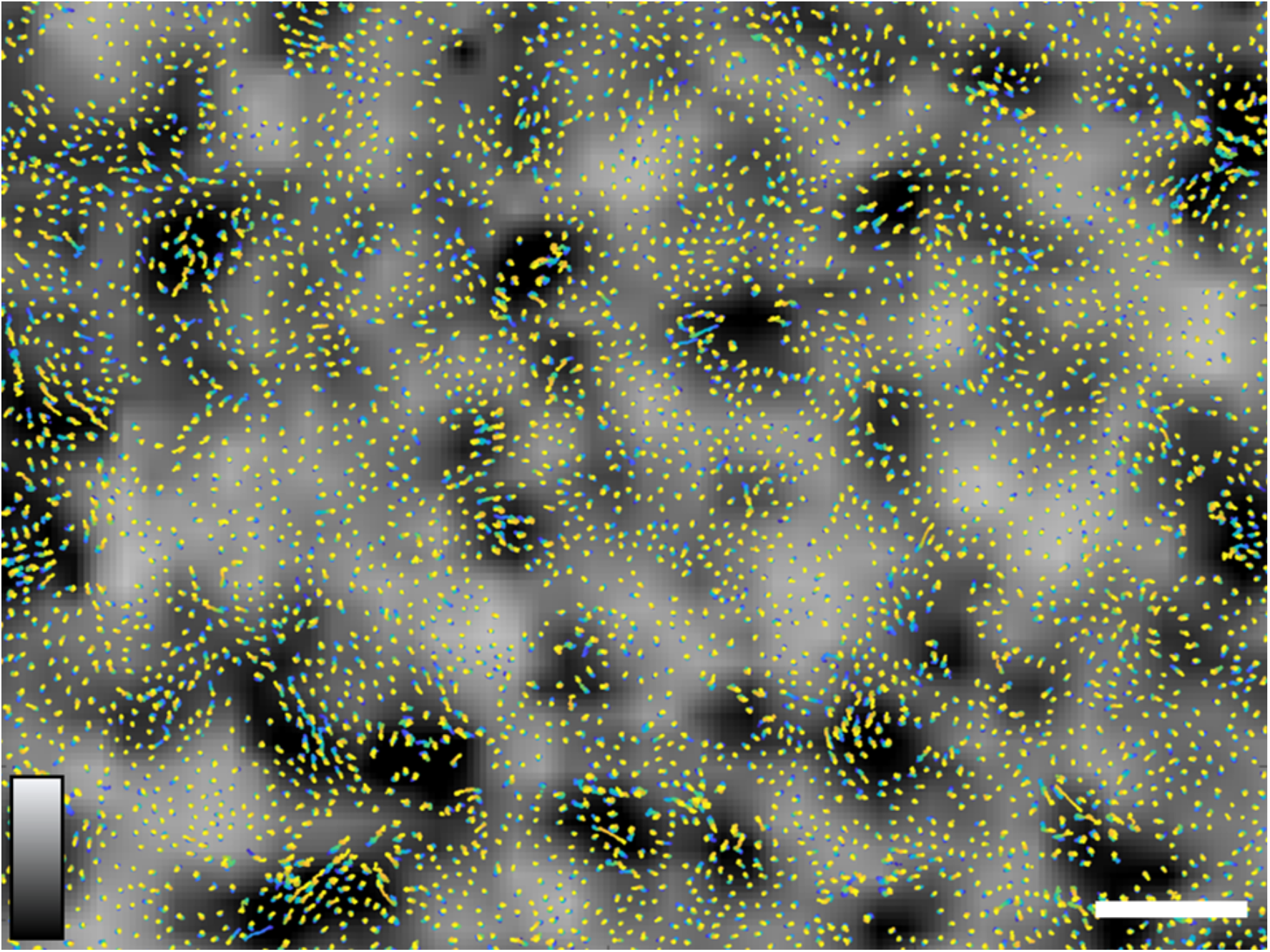
Gray map of self-overlap parameter *Q*_*i*_(*a*, *δt*) where *a* = 1.2 μm is the reference length scale, *δt* = 200 min is the time delay, along with cell trajectories from *t*=0 (dark blue) to *t*=1625 min (light yellow). *Q*_*i*_(*a*, *δt*) decays exponentially from 1 to 0 as cells gradually move away from their initial reference positions. There are regions of different sizes with small and large values of *Q*_*i*_(*a*, *δt*), showing the dynamic heterogeneity of the system. The color bar scales linearly from 0.5 (black) to 1 (white). The scale bar is 100 μm.

**Fig. S6.**
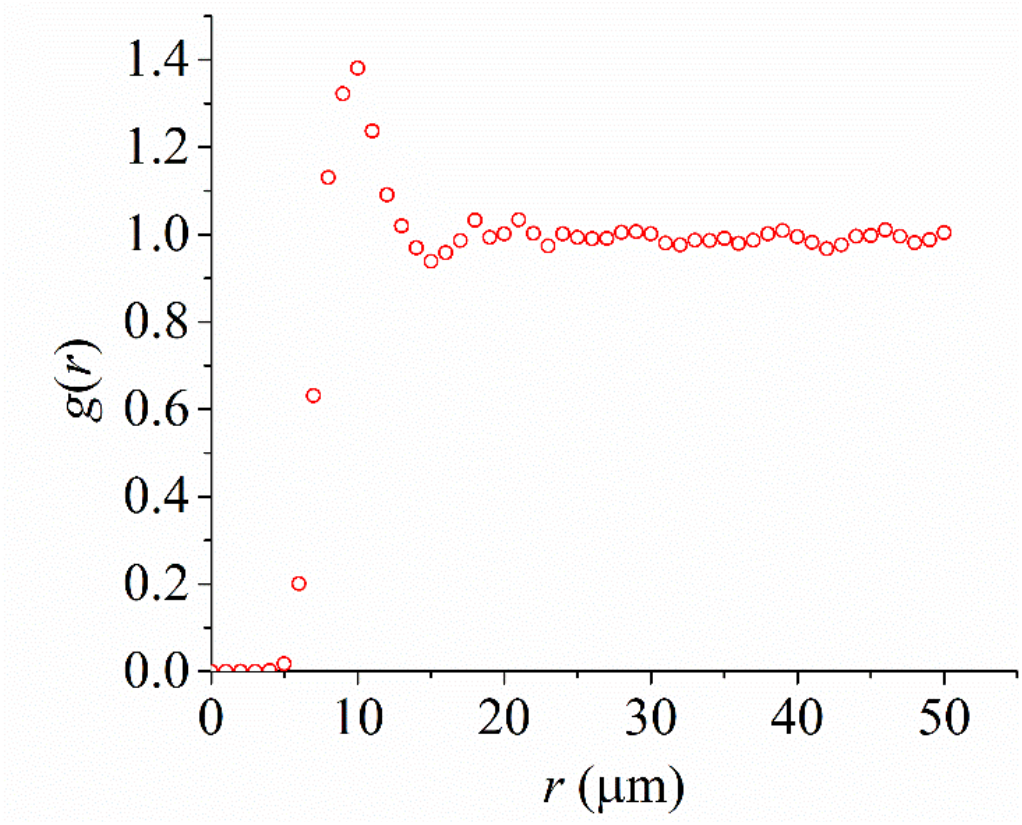
1D radial distribution function of the epithelial tissue averaged radially in space. The first valley gives a threshold distance *r*^*^ = 15 μm.

**Fig. S7.**
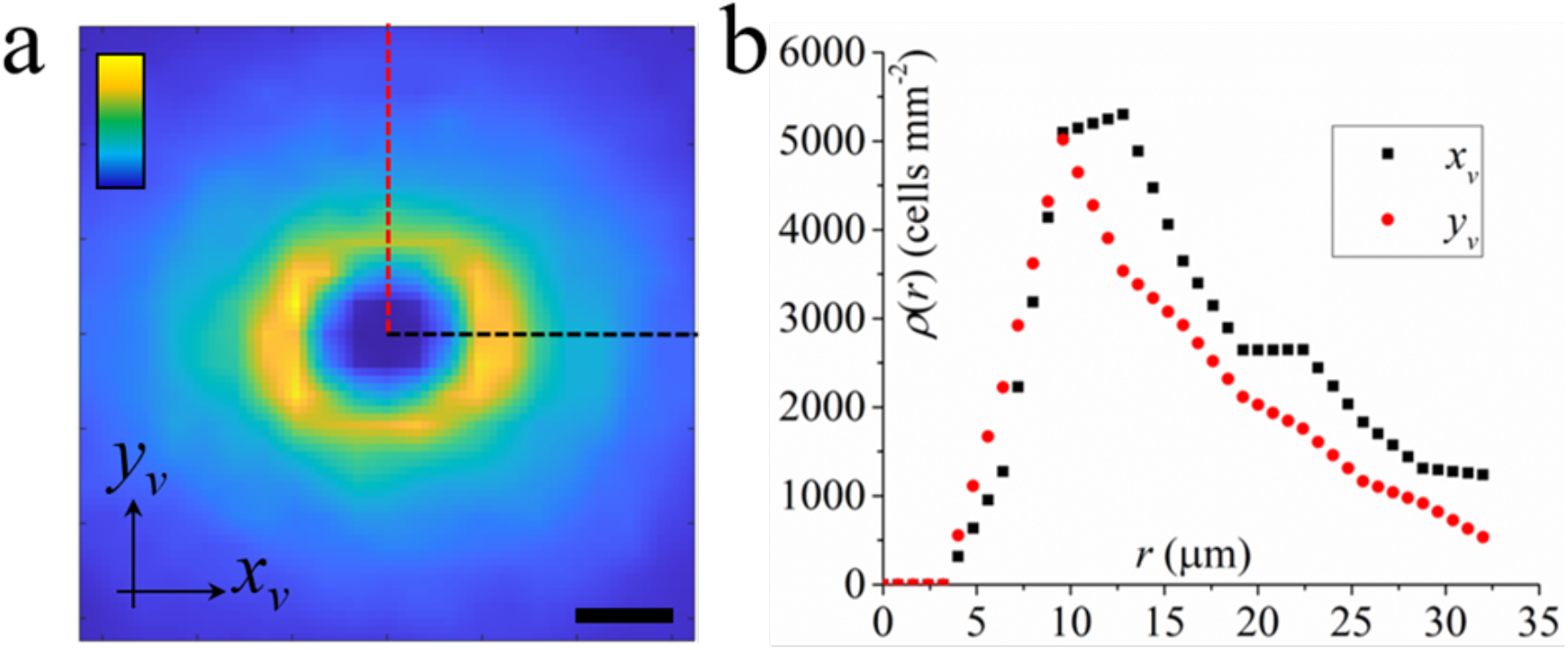
(a) 2D radial distribution function of cells in large fast cellular clusters in a local coordinate system where the velocity of the reference cell is always aligned along the positive direction of the *x*_*v*_-axis. The function is not normalized by the mean cell density of the sample, thus the color bar represents cell density which scales linearly from 0 to 6000 (cells per mm^2^). Scale bar is 10 μm. (b) The 1D cross profiles of the 2D function along the *x*_*v*_-axis (black dashed line) and *y*_*v*_-axis (red dashed line), respectively.

**Fig. S8.**
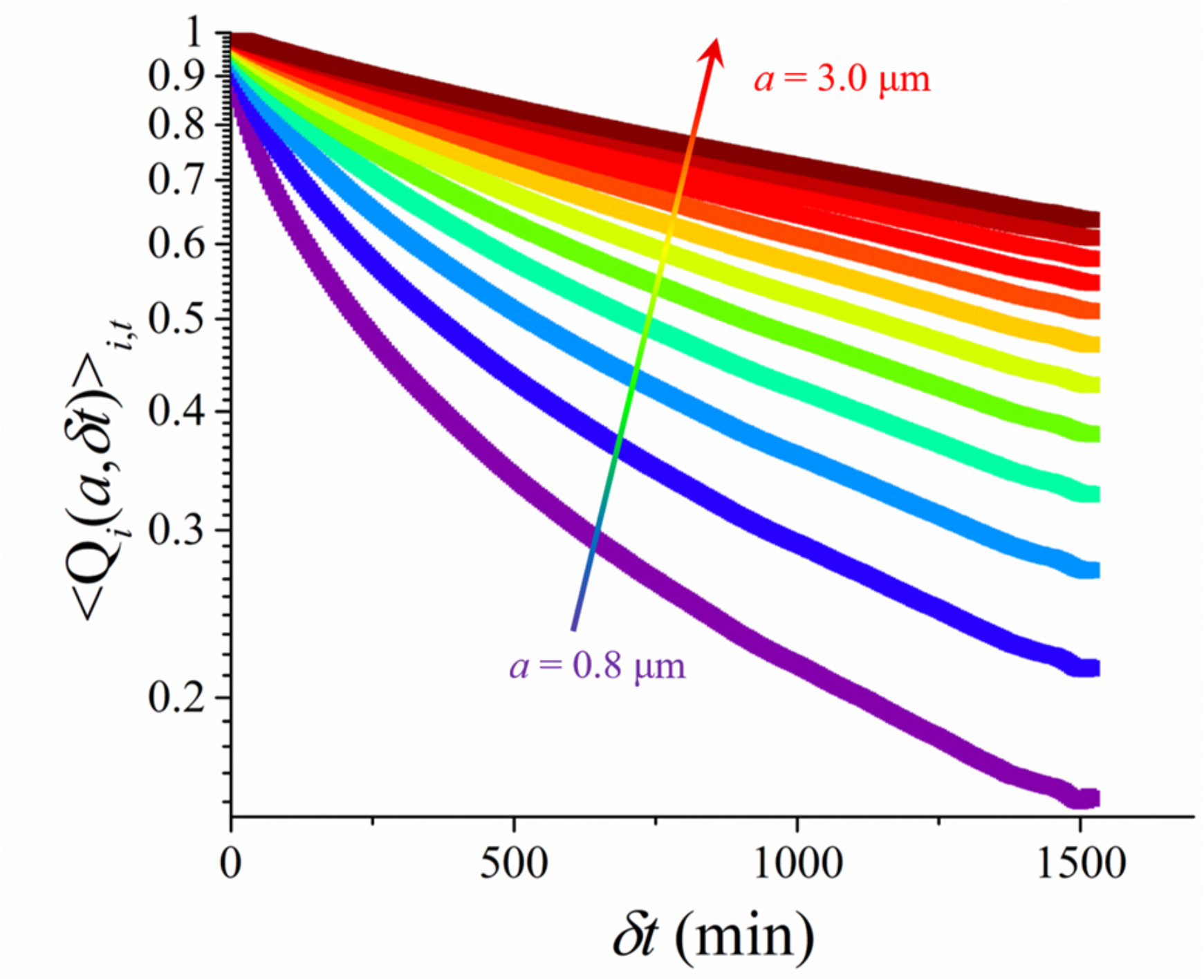
The temporal dependence of the overlap function. The reference length, *a*, increases from 0.8 to 3.0 μm in a step of ∼0.2 μm as indicated by the arrow.

**Fig. S9.**
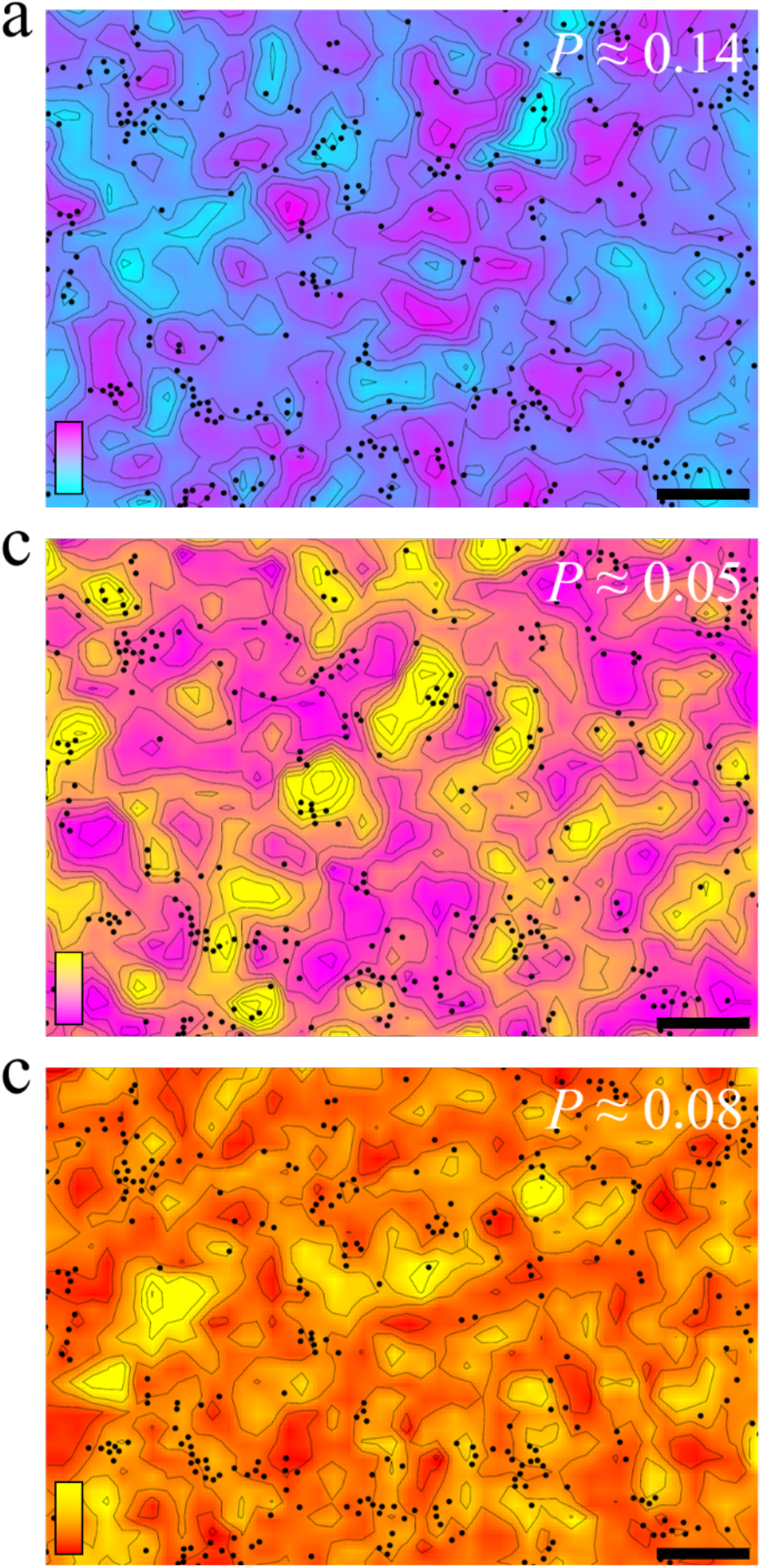
Contour maps of local density *ρ* (a), local nematic order *O*^2^ (b), and local hexatic bond orientational order *O*^6^ (c). The black dots represent the cells with the 10% smallest values of self-overlap parameter. The color bars scale linearly from 6×10^-3^ to 13×10^-3^ for *ρ*, from 0.3 to 0.6 for *O*^2^ and from 0.3 to 0.5 for *O*^6^. Scale bars are 100 μm. *P* gives the Pearson correlation coefficient and one recognizes that the correlations are weak.

